# A draft *de novo* assembly of *Diadema antillarum*, a keystone herbivore of the Caribbean reefs

**DOI:** 10.64898/2026.05.24.727502

**Authors:** Audrey J. Majeske, Juliet M. Wong, Carlos Farkas Pool, Jose Eirin-Lopez, Walter Wolfsberger, Nikolaos V. Schizas, Alondra M. Díaz-Lameiro, Stephanie O. Castro-Márquez, Kenneth Hilkert, Alejandro J. Mercado Capote, Tarás K. Oleksyk

**Affiliations:** Oakland University, MI, USA; University of Puerto Rico at Mayagüez, PR, USA; Duke University, NC, USA; Florida International University, FL, USA; Universidad Católica de la Santísima Concepción, Biobío Region, Chile

**Keywords:** long-spined black sea urchin, genome assembly, marine invertebrate, Puerto Rico

## Abstract

We generated the first reference-level nuclear genome assembly of the keystone Caribbean long-spined black sea urchin species, *Diadema antillarum* (Philippi, 1845). Using whole-genome sequencing data from PacBio HiFi, Oxford Nanopore, and Illumina platforms, we employed multiple assembly strategies to generate a high-quality, near-complete genome. The final assembly spans 1.73 Gbp, consists of 2,964 scaffolds, and has an N50 of 1.56 Mbp. BUSCO analysis (metazoa_odb10) indicates 98.4% completeness. The genome displays a heterozygosity rate of 2.52% and contains 42.85% repetitive elements, of which 29.96% are unclassified. Coverage analysis reveals that while most of the genome was assembled at 11x depth, certain regions exhibit up to 530x coverage. Notably, regions exceeding 33x coverage account for 30.53% of the repetitive content, suggesting localized expansion of repeats. Duplication analysis of the assembled contigs shows that approximately 66% of contigs have duplicated, which supports segmental genome duplication in the past, and is further evidenced by the moderate level of heterozygosity of the assembly. While these characteristics contribute to the complexity of the genome, they do not diminish the quality of our assembly. Despite this complexity, our assembly maintains high completeness and contiguity. Our assembly provides a valuable resource for future genetic studies and serves as a critical framework for conservation, monitoring, and restoration of *D. antillarum* populations across the Caribbean.

## Introduction

The long-spined black sea urchin, *Diadema antillarum* is a keystone herbivore within Caribbean coral reefs, whose presence has been directly linked to improved coral settlement, growth, and survivorship (Edmunds & Carpenter, 2001; Idjadi et al., 2010). Historically abundant, in the early 1980s, *D. antillarum* suffered a mass mortality event that removed an average of 98.06% of *D. antillarum* populations across the Caribbean, contributing to the considerable shifts from coral-to algae-dominated reefs (Lessios, 2016). Over the next few decades, *D. antillarum* slowly recovered (Lessios, 2016), despite several smaller scale mortality events (Forcucci, 1994; Lessios, 1988). However, in 2022 another die-off event occurred across many regions in the Caribbean (Hylkema et al., 2023; Levitan et al., 2023), later reported to have been caused by a scuticociliate similar to *Philaster apodigitiformis* (Hewson et al., 2023). Mass mortalities due to disease outbreaks are predicted to increase in frequency as climate change and other anthropogenic stressors continue (Harvell et al., 1999). As such, more die-offs of *D. antillarum* are expected to occur in the future, which will lead to further degradation of Caribbean coral reefs and potential further loss of genetic diversity within the species.

Investigations into these mass mortalities in *D. antillarum* continue, including efforts to understand the mechanisms of transmission and infection, as well as determine the many population- and ecosystem-level impacts. Additionally, many efforts have focused on *D. antillarum* conservation and restoration as a means of combating die-offs and augmenting natural population recovery. For instance, methods have been developed for the aquaculture of *D. antillarum* for restocking and population enhancement (Hassan et al., 2022; Pilnick et al., 2021, 2022; Wijers et al., 2023), juveniles have been reintroduced into reefs (Williams, 2018, 2022), artificial settlement substrates have been deployed to increase densities of recruits (Hylkema et al., 2023), and artificial shelters and habitat structures have been constructed to reduce loss from predation (Dame, 2008; Delgado & Sharp, 2021). Overall, the substantial ecological value, yet vulnerability, of this species is well-recognized and as such, considerable amounts of time, money, and resources have been put towards *D. antillarum* research.

Given their beneficial ecological role within Caribbean coral reefs, history of extensive mortality events, and the sizable efforts invested in assisting their recovery, surprisingly few molecular resources are available for *D. antillarum*, although its mitochondrial genome was recently completed (Majeske et al., 2022). That work corrected inaccuracies in family-level phylogenetic relationships within Diadematidae based on the four available mitochondrial genomes sequenced to date (Majeske et al., 2022). Nuclear genomes, however, provide a far more comprehensive framework for comparative analyses and can further resolve these relationships. Nuclear genome assemblies are currently available for 24 separate echinoids in the NCBI database, including one within Diadematidae, namely *Diadema setosum* (Hong Kong Biodiversity Genomics Consortium, 2024). Nonetheless, this set of resources have revealed that echinoids possess unusually expanded and diversified innate-immune gene families, including Toll-like receptors (TLRs), NACHT- and leucine-rich repeat–containing receptors (NLR/NACHT-LRR), scavenger receptor cysteine-rich proteins (SRCR), and the rapidly evolving Trf/Sp185/333 family (Polinski et al., 2024; Smith et al., 2018). Here we present the first reference-level nuclear genome assembly for *D. antillarum*. We also begin to explore genomic diversity between *D. setosum* and *D. antillarum* assemblies (Supplemental Table 1). This additional resource will be instrumental for future more thorough comparative immunological and population genetic studies and will also aid in reef monitoring and recovery efforts for this key marine invertebrate species.

**Table 1.**
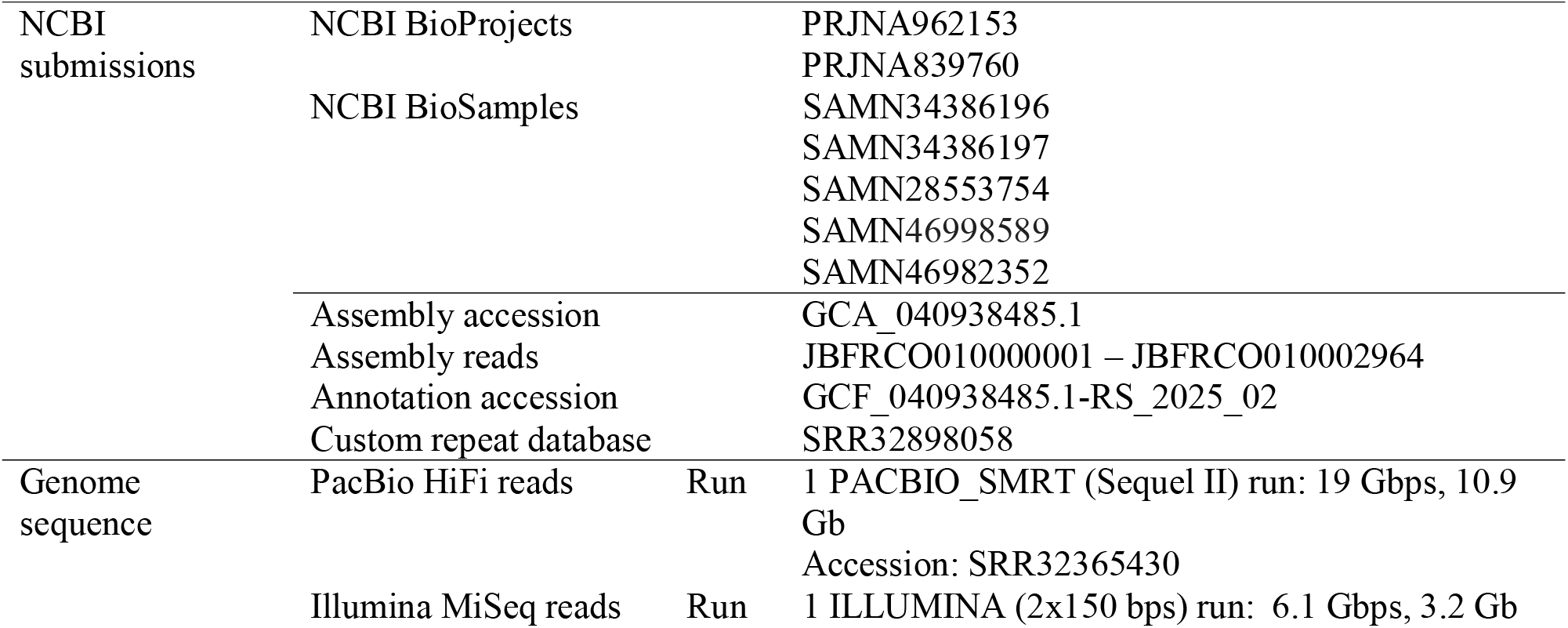
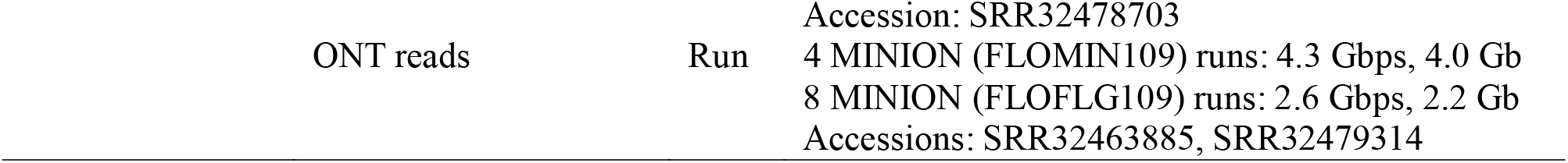
NCBI accession numbers and genome sequences.

## Methods

### Biological materials

Two sea urchins (An #1 and An #2) were collected in Rincón, Puerto Rico in 2018 (18°20′35.2′′N, 67°15′40.5′′W; Figure 1). For the An #1 specimen, whole coelomic fluid (wCF) containing coelomocytes was withdrawn through the peristomal membrane using a sterile 23-gauge needle connected to a 5-mL syringe. The animal was photographed and then returned to its habitat at the collection site (Figure 1). Duplicate sample tubes (containing 1 mL each) were held on ice during transport to the University of Puerto Rico Mayagüez (UPRM). Next, the tubes were centrifuged to pellet the coelomocytes, and the cell free fluid was removed from the pellet which was replaced with RNAlater solution (Invitrogen) to amply cover each pellet. Samples were stored frozen before shipment on ice packs to Oakland University (OU), Rochester MI, followed by frozen storage.

**Figure 1.**
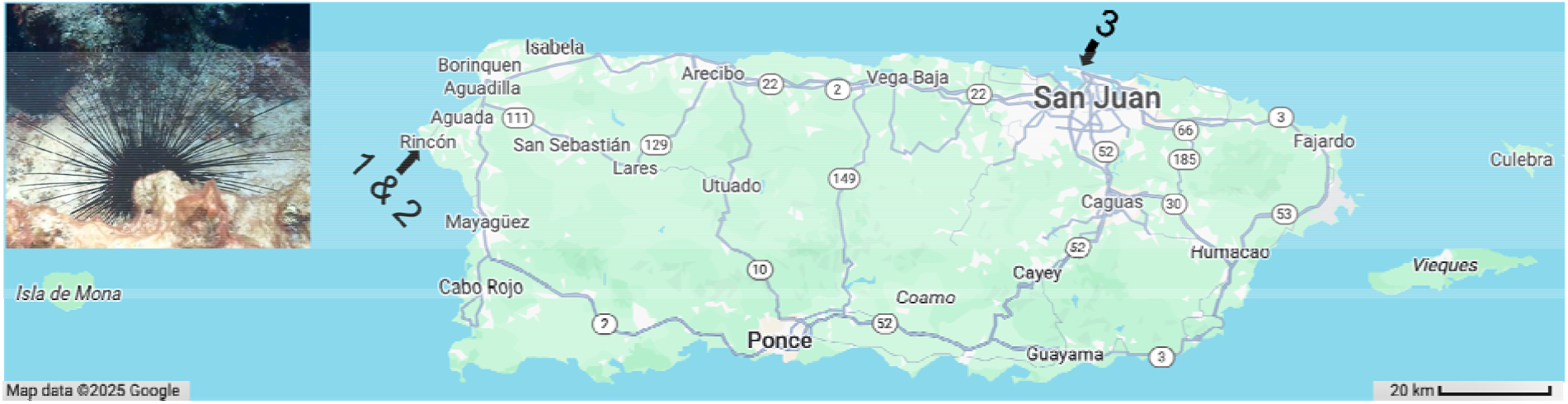
Map of Puerto Rico and collection sites of adult Diadema antillarum. Animals #1 and #2 were collected in Rincon (18°20′35.2′′N, 67°15′40.5′′W) in 2018. Animal #3 was collected in Punta Escambrón (18° 27’ 59.70” N, 66° 5’ 10.26” W) in 2022. Map source: Modified from Google Maps (accessed February 14, 2025). Inset photo of an D. antillarum in its natural habitat taken by A. J. Mercado Capote.

The second adult sea urchin (An #2) was transported live to the lab at UPRM. There, the Aristotle’s Lantern was carefully dissected and removed. The animal then was inverted to decant whole coelomic fluid into a sterile 50 mL conical tube. Approximately 15 mL of wCF was collected from the animal. The wCF was aliquoted into microcentrifuge tubes (1 mL per tube) and coelomocytes were processed in the same manner as An #1. All samples were stored in a -20 °C freezer prior to shipment on ice packs to OU. The samples from both animals were stored frozen at OU prior to DNA extraction.

A third sea urchin was collected (An #3) at Punta Escambrón, Puerto Rico (18° 27’ 59.70” N, 66° 5’ 10.26” W; Figure 1) on May 11, 2022, by SCUBA. The animal was immediately transported to facilities at the University of Puerto Rico at Río Piedras, where it was spawned by intracoelomic injection of 0.53M potassium chloride. Sperm was collected “dry” by pipetting it directly from the test. The sperm was then rinsed three times using a 1X solution of molecular biology grade phosphate buffered saline (PBS). Sperm cells were then concentrated by centrifugation and excess PBS was removed via pipetting. The sperm sample was frozen at -20 °C and transported to facilities at Florida International University prior to DNA isolation.

### Library preparation and sequencing

For the An #1 and An #2 coelomocyte samples, the RNAlater solution was removed prior to performing a high molecular weight (HMW) DNA extraction method using a Monarch HMW DNA Extraction Kit (New England Biolabs) according to the manufacturer’s instructions. Sample quality and quantity was assessed using an Implem C40 NanoPhotometer. DNA samples from both animals were prepared for long read sequencing with a SQK-LSK109 ligation sequencing kit (Oxford Nanopore Technologies; ONT). Library quantities were examined on a Qubit 2.0 fluorometer after sample preparation using a Qubit dsDNA HS Assay Kit (Thermo Fisher Scientific). Samples were sequenced separately which culminated in eight different Flongle (R9.4.1 or R10.4.1) and four different MinION (R9.4.1 or R10.4.1) flow cells using a MinION sequencing system.

The DNA extraction from sperm (An #3) was performed using an E.Z.N.A. DNA/RNA Isolation Kit (Omega Bio-Tek). Quality control was performed using a NanoVue Plus Spectrophotometer (GE), a Qubit 4 Fluorometer (Invitrogen), and TapeStation (Agilent) to verify suitable quantity, purity, and integrity of the DNA sample. Total genomic DNA was submitted to the University of Florida’s Interdisciplinary Center for Biotechnology Research (UF ICBR) for SMRT bell library construction and PacBio HiFi sequencing. The prepared sample was sequenced on a SEQUEL IIe SMRT cell with diffusion loading, 2-hr pre-extension, on-instrument HiFi reads generation, and 30 hr movies. An aliquot of the sperm DNA sample was also used for standard Illumina library construction (New England Biolabs) and sequenced on an Illumina MiSeq 2×150 cycles at UF ICBR. All libraries were analyzed for quality and quantity prior to sequencing.

ONT reads were generated using Guppy (v6.1.5), which included automated base-calling a post-processing stages to remove adaptor sequences and retain high quality reads with a Phred score of ≥ 30. The data files from multiple ONT runs were combined for An #1 and An #2 prior to downstream assembly applications. Sequences obtained using an Illumina platform were assessed for read quality using FastQC. Adaptor sequences were removed using Trimmomatic. Sequences were deposited into the NCBI database (Table 1).

### Genome assembly and annotation

The full list of the relevant software tools and versions used in this project are included in Table 2. First, adaptor sequences of HiFi reads were removed using HiFiAdapterFilt. Then, the adaptor free reads were run through Jellyfish for kmer counting of size 21 and the output file was assessed for estimations of genome size, heterozygosity and repetitiveness using Genomescope2.0. In addition, ploidy was assessed using Smudgeplot (v0.2.6). In this process, kmer counting of size 21 and 31 for the adaptor removed HiFi reads was completed using FastK (v1.1), and the output file was used to generate the Smudgeplot.

**Table 2.**
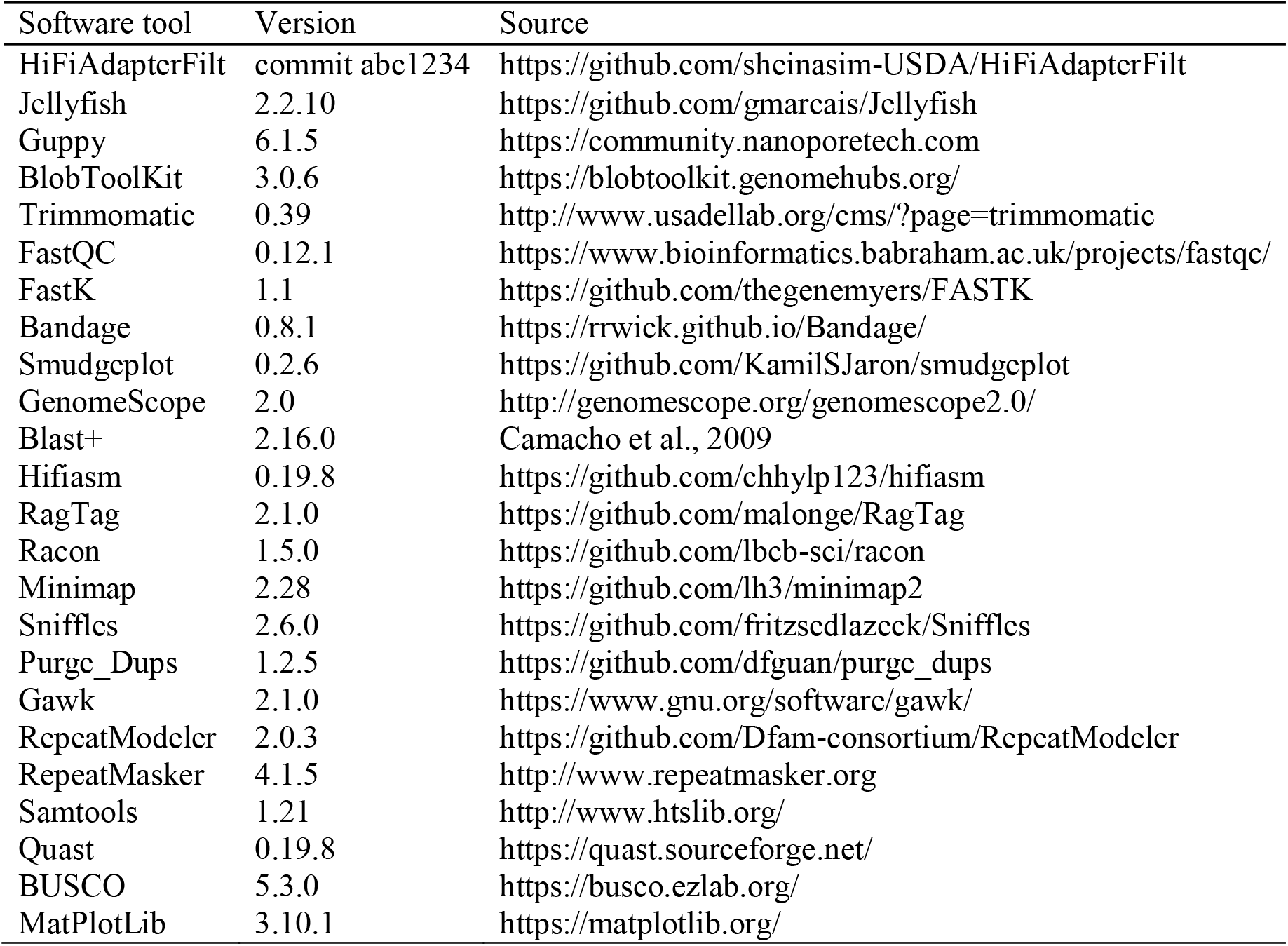
Assembly and annotation software usage.

To maximize the quality of the final genome assembly, we employed different assembly pipelines (Table 3) and then compared the output assembly results for contiguity, completeness, and quality metrics. Contamination screening for the presence of other diploid DNA or other prokaryotic species DNA was completed for all reads generated by the different platforms using Blast+ 2.16.0 (Camacho et al., 2009). All pipelines used the Hifiasm (v0.19.8) in the default haploid mode. In the first pipeline (*HiFiasm v0*.*1*), HiFi reads were assembled with no additional scaffolding, polishing, or error correction applied. This assembly served as a primary reference and baseline for comparison across all workflows. In a second pipeline (*HiFiasm v0*.*2*), Hifiasm was used to simultaneously assemble HiFi and ONT reads. This hybrid assembly was performed without additional polishing or structural refinement steps. Following initial assembly, RagTag (v2.1.0) was implemented to improve contiguity and address missing sequence regions. In one workflow (*HiFiasm v0*.*3*), RagTag was used to contig a HiFi assembly using ONT Hifiasm assembled reads as a reference for structural guidance to improve the order and orientation of the contigs. In the last pipeline (*HiFiasm v0*.*4*), which served as the final assembly, HiFi reads were first assembled with Hifiasm, followed by RagTag patching using the assembled ONT reads, and then polishing with Illumina short reads using Racon.

**Table 3.** Tools used for different genome assembly pipelines.

| Assembly version | Assembled Reads | Post assembly processing |  |
| --- | --- | --- | --- |
|  |  | Ragtag Scaffolding | Racon Polishing |
| *Hifiasm v0.1* | HiFi | – | – |
| *Hifiasm v0.2* | HiFi + ONT | – | – |
| *Hifiasm v0.3* | HiFi | ONT reads | – |
| *Hifiasm v0.4* | HiFi | ONT reads | Illumina reads |

Next, we used BlobToolkit to generate a snail plot of the *HiFiasm v0*.*4* assembly. We used Quast (v5.2.0) without a reference genome to assess the quality of the final assembly. We used Bandage (v0.8.1) to visualize the assembly. We used Minimap (v2.28) to align the raw HiFi reads back to the assembly and then converted the output to a .txt file using Samtools (v1.21). We generated a plot of the coverage statistics using MatPlotLib (v3.10.1). We used GNU Awk (Gawk v5.1.0) to output coverage statistics from the mapped reads including mean, median, minimum, maximum coverage, the number of total raw bases, mapped bases, unmapped bases, bases with no coverage/missing bases, and highly variable/soft-clipped bases (using the 10S100M CIGAR string). We also calculated the percent genome coverage by all bases, using percent_coverage=$(echo “scale=5; ($total_bases_with_coverage / $genome_length) * 100” | bc). We extracted the high coverage reads (≥ 100x) as well as those encompassing three times the median read coverage (11x), or ≥ 33x. This was accomplished by first extracting the high coverage bases using awk ‘$3 > 100 {print $0}’, or the ≥ 33x coverage reads using awk ‘$3 > 33 {print $1}’ and then converting each .txt file into a .bed file using awk ‘{OFS=“\t”; print $1, $2-1, $3}’. To extract the mapped reads with the specified coverage (≥ 100x or ≥ 33x), we converted the .bed file to a .bam file using samtools view. Each .bed file was then converted to a fasta file via samtools fasta, which outputted the mapped high coverage reads or those with ≥ 33x coverage. We also extracted large structural variants from the HiFi reads using Sniffles (v2.6.0). We filtered these variants by type (SVTYPE=DEL, INS, DUP) and size (length >= 1000 bp). Then we counted and outputted the number of deletions, insertions and duplications as well as the total bps associated with the mapped reads for each type of structural variant. Next, we extracted the large structural variant duplicated reads from the .vcf file using grep -E “SVTYPE=DUP” followed by awk -F’\t’ ‘{split($8,a,”;”); for (i in a) if (a[i] ∼ /SVLEN/) print $1, $2, $3, a[i]}’ to output a text file. We converted this text file to a .sam file with Samtools (v1.21) and then aligned the extrapolated reads back to the assembly using Minimap (v2.28). We then generated a histogram of the distribution of read length and depth of coverage for these reads.

We used RepeatModeler (v2.0.3) to generate a custom repeat database of *D. antillarum*. This database was used to characterize repeats in the final assembly using RepeatMasker (v4.1.5). We also used this custom repeat database to map back the previously extracted high coverage or the ≥ 33x coverage read files as a query using RepeatMasker (v4.1.5). Additionally, we used Purge_Dups (v1.2.5) to assess the contig redundancy of the assembly against the custom repeat database. The cutoff values used by the Purge_Dups tool were 5, 9, 15, 18, 30, 54, which were specific to the depth of coverage analysis results of the assembly. We extracted the number of contigs in each of the two output files (named hap. fa and purged. fa) using grep -c “^>“ hap. fa. We also extrapolated the total bps in the two output files using awk ‘/^>/ {if (seqlen){sum+=seqlen}; seqlen=0; next} {seqlen += length($0)}. We performed a BUSCO analysis to examine the results of the ‘cleaned’ assembly output following the Purge_Dups analysis. We used Minimap (v2.28) to align the two output files using the purged.fa (‘cleaned’ assembly) as a reference and retrieved the output mapped file, which was then used in a custom python script to report statistical results (https://doi.org/10.5281/zenodo.15172401).

The *HiFiasm v0*.*4* assembly was annotated by NCBI using the Eukaryotic Genome Annotation Pipeline (EGAP). The automated pipeline was used to integrate transcript and protein alignments from RefSeq (if available), *ab initio* gene prediction models, repeat masking, and RNA-seq data to produce high-confidence gene models and functional annotations. The pipeline assigned gene symbols, product names, and GO terms based on curated databases.

## Results

### Data Description

The file sizes and total bps generated for the samples using ONT or Illumina are listed in Table 1. PacBio yielded 2,878,999 reads containing 19,633,500,452 bps. The mean raw read length was 6,819 bps. The mean HiFi read quality was Q34 and the mean number of sequencing passes was 13. Results of Genomescope2.0 analysis predicted a haploid genome length of 703,182,906 bps, with 76.7% uniqueness, a heterozygosity of 2.52%, a duplication ratio of 0.582%, and an error rate of 0.195% (Figure 2A). The model fit accuracy was 97.75 ± 1.17%. Results of the Smudgeplot for kmer size 21 include four genotype configurations AB, AAB, AAAB, and AABB with the respective frequencies 0.61, 0.23, 0.03, and 0.05. For kmer 31, the Smudgeplot depicts only two genotypes, AB and AAB, with frequencies of 0.73 and 0.18, respectively (see Supplementary Figure 1).

**Figure 2.**
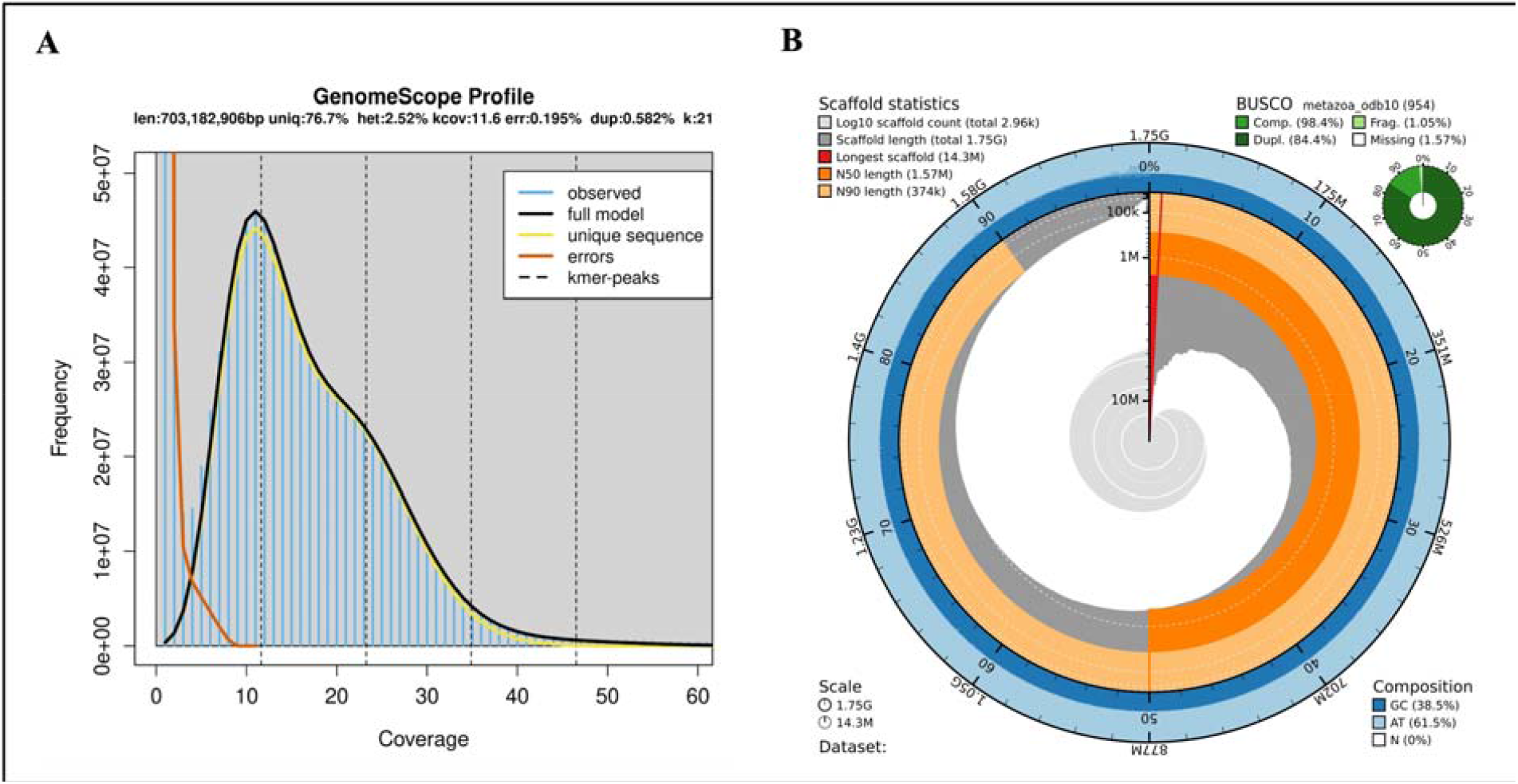
Quality metrics for the HiFi reads and assembly statistics for the final *Diadema antillarum* genome assembly (*HiFiasm v0*.*4*). A. K-mer spectrum output of data without adaptors using GenomeScope2.0. The k-mer distribution is unimodal, with a primary peak at ∼11x coverage, consistent with haploid genome content. A minor shoulder around ∼25x likely reflects high-copy-number or repeated sequences (Vurture et al. 2020). B. BlobToolKit Snail plot showing a graphical description of the quality metrics in Table 4 for the *HiFiasm v0*.*4* assembly. The plot circle represents the full size of the assembly. The red line represents the size of the longest scaffold, and all other scaffolds are drawn in dark gray and arranged in size-order moving from a clockwise direction around the circular plot starting from the outside of the central position. Dark orange shows the N50 value and light orange shows the N90 value. The central light gray spiral represents the cumulative scaffold count with the white line at each order of magnitude. The white regions in this area indicate the proportions of Ns in the assembly. The outer dark and light blue ring shows the mean, maximum, and minimum GC vs. AT content at 0.1% intervals. A summary of complete, fragmented, duplicated, and missing BUSCO genes in the metazoan_odb10 set is shown in the top right (Challis et al. 2020).

### Genome assembly and annotation

Table 4 lists the quality metrics of the different assembly pipelines. The final assembly (*HiFiasm v0*.*4*) was composed of 2,964 scaffolds, with a length of 1,754,675,526 bps (1.7 Gbps). The largest scaffold of the assembly was (14,326,224) 14.3 Mbps, the N50 was 1.56 Mbps, the N90 was 374 Kbps, and the GC level was 38.48%. Similar results are also depicted in the snail plot (Figure 2B). The BUSCO analysis for the assembly indicated a completeness of 98.4% based on the metazoan database (metazoan_odb10) containing 954 gene families. Among the complete BUSCOs, 14.0% were single-copy, while 84.4% were duplicated. In addition, 1.0% were fragmented BUSCOs, and 0.06% were missing BUSCOs. The Quast results of the final assembly calculated a duplication ratio of 2.429.

**Table 4.**
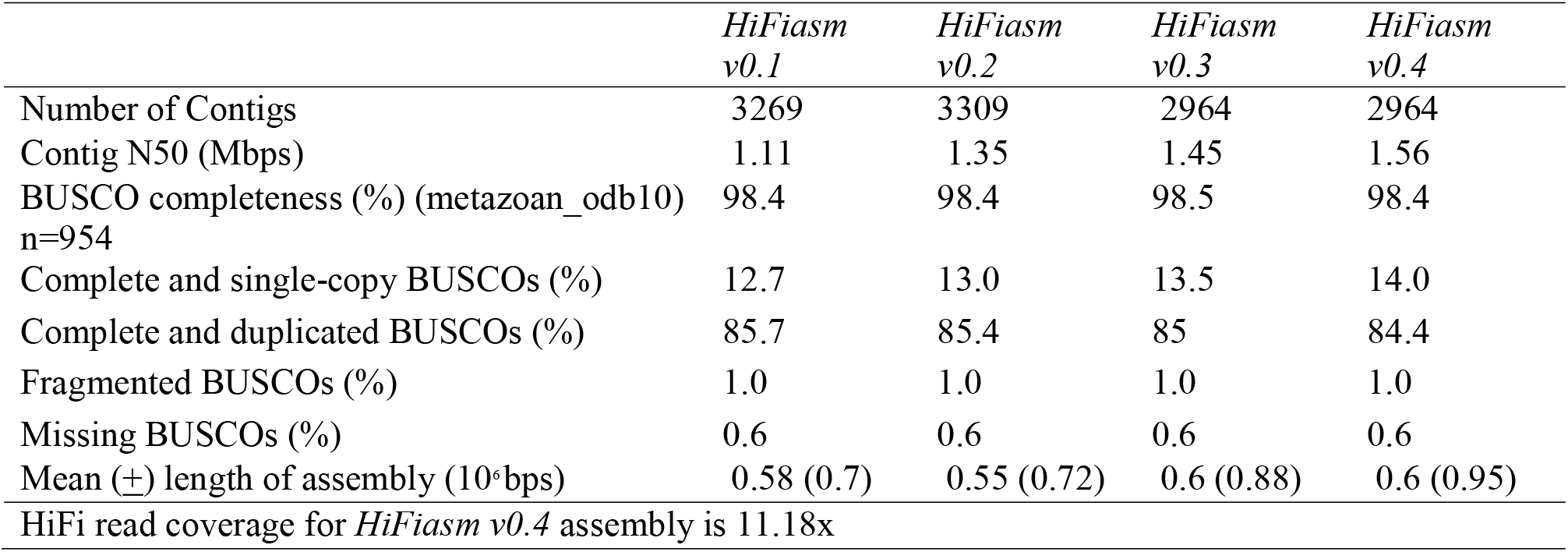
Metrics for genome assembly pipelines.

|  | *HiFiasm*<br>v0.1 | *HiFiasm*<br>v0.2 | *HiFiasm*<br>v0.3 | *HiFiasm*<br>v0.4 |
| --- | --- | --- | --- | --- |
| Number of Contigs | 3269 | 3309 | 2964 | 2964 |
| Contig N50 (Mbps) | 1.11 | 1.35 | 1.45 | 1.56 |
| BUSCO completeness (%) (metazoan_odb10)<br>n=954 | 98.4 | 98.4 | 98.5 | 98.4 |
| Complete and single-copy BUSCOs (%) | 12.7 | 13.0 | 13.5 | 14.0 |
| Complete and duplicated BUSCOs (%) | 85.7 | 85.4 | 85 | 84.4 |
| Fragmented BUSCOs (%) | 1.0 | 1.0 | 1.0 | 1.0 |
| Missing BUSCOs (%) | 0.6 | 0.6 | 0.6 | 0.6 |
| Mean (±) length of assembly ($10^6$bps) | 0.58 (0.7) | 0.55 (0.72) | 0.6 (0.88) | 0.6 (0.95) |
| HiFi read coverage for *HiFiasm* v0.4 assembly is 11.18x |  |  |  |  |

The base pair coverage plot of the *HiFiasm v0*.*4* assembly is shown in Figure 3. The initial peak in the histogram indicates that most of the genome is covered at a depth of 11x, however the long tail includes per base sequencing depths with a much higher coverage (from 100x up to 530x). Other coverage statistics indicated the number of mapped bases was 19,707,975,605, which was 74,475,153 bps more than the total number of raw bases, or 0.38% of the reads had mapped more than once. There were zero unmapped bases, the total number of no coverage bps was 1,443,017,540, and the total number of highly variable/soft-clipped bps was 543,248,118. The Bandage plot of the long read assembly depicted individual lines, each representing one of the 2,964 contigs, with no connecting edges between the contigs. The longest contig in the plot contained 14,326,224 bps, and the shortest contig contained 12,568 bps (see Supplementary Figure 2).

**Figure 3.**
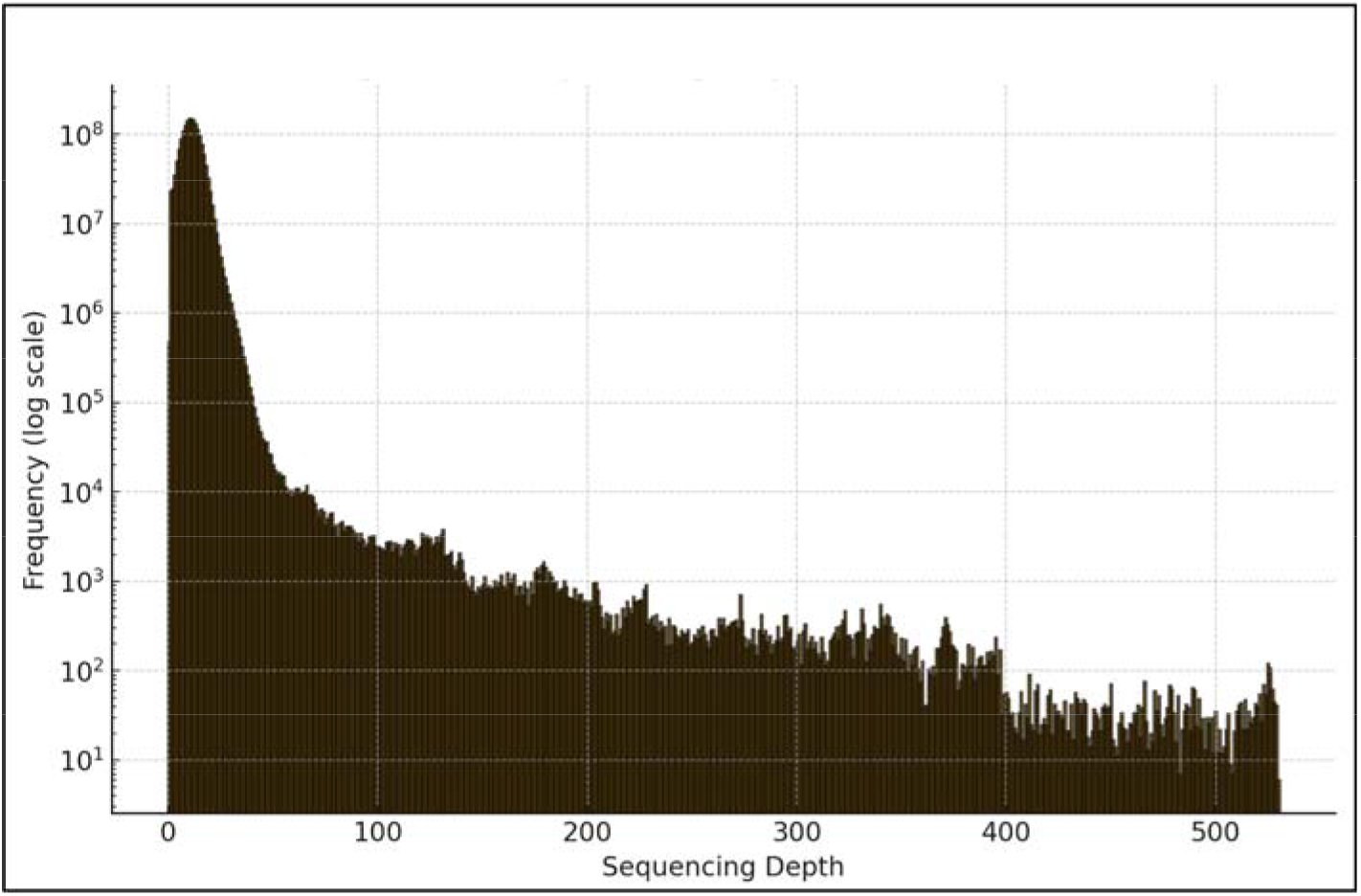
Histogram of HiFi sequencing depth across the *HiFiasm* v0.4 genome assembly of *Diadema antillarum*. HiFi reads were mapped back to the assembly using Minimap (v2.28), and then per-base coverage statistics was performed to summarize the sequencing depth across the genome. The histogram displays the frequency distribution of sequencing depth on a logarithmic scale and was generated using Matplotlib (v3.10.1). The primary peak at 11x coverage indicates that the majority of the genome is represented at this depth, with minimal regions lacking coverage, suggesting a highly complete and representative assembly. A long right-skewed tail extends from the main peak, with localized regions showing substantially higher coverage, ranging from 100x to 530x, likely corresponding to high-copy or repetitive elements within the genome (Miller et al. 2019).

Regarding the output statistics of the large structural variant analysis, there were 1,317 separate deletions comprising a total of 4,312,104 bps (4.3 Mbps) in the genome, in which the length of the longest single deletion was 119,558 bps. For insertions, there were 431 individual insertions totaling 820,154 bps. While there were only 31 separate duplicated reads, they comprised a total of 5,406,802 bps (5.4 Mbps), with an average length of 174,413 bps for a single duplication, and the longest duplication spanned 2,449,537 bps (2.4 Mbps). The plot showing the distribution of read length and the coverage plot for the large structural duplications (≥ 1000 bps) mapped back to the assembly are shown in Supplementary Figure 3. In Supplementary Figure 3A, most of the large, duplicated reads fall between 6,000 to 10,000 bps in length, which indicates long read consistency at these high coverage regions. The histogram plot of the large, duplicated reads shows a consistent range of coverage up to 80x that is devoid of major spikes (see Supplementary Figure 3B).

Table 5 summarizes the RepeatMasker results for the *HiFiasm v0*.*4* assembly, the extracted ≥ 33x coverage reads, and the high coverage reads. In short, for the assembly, the total percentage of repeat elements was 42.84% (751,856,782 bps), of which 29.96% were unclassified. The total length for ≥ 33x coverage reads was 373,089,082 bps, which represented 21.2% of the assembly. For these reads, 61.52% contained repeat elements (229,538,485 bps). Thus, the repeats with ≥ 33x read coverage of the genome comprise 30.53% of the total repeat elements in the assembly. The total bps for the high coverage (≥ 100x) reads was 49,529,407, which represented 2.82% of the assembly. For the high coverage reads, 93.07% contained repeat elements. Therefore, the reads with ≥ 100x coverage of the genome comprises 6.59% of the total repeats in the assembly.

**Table 5.**
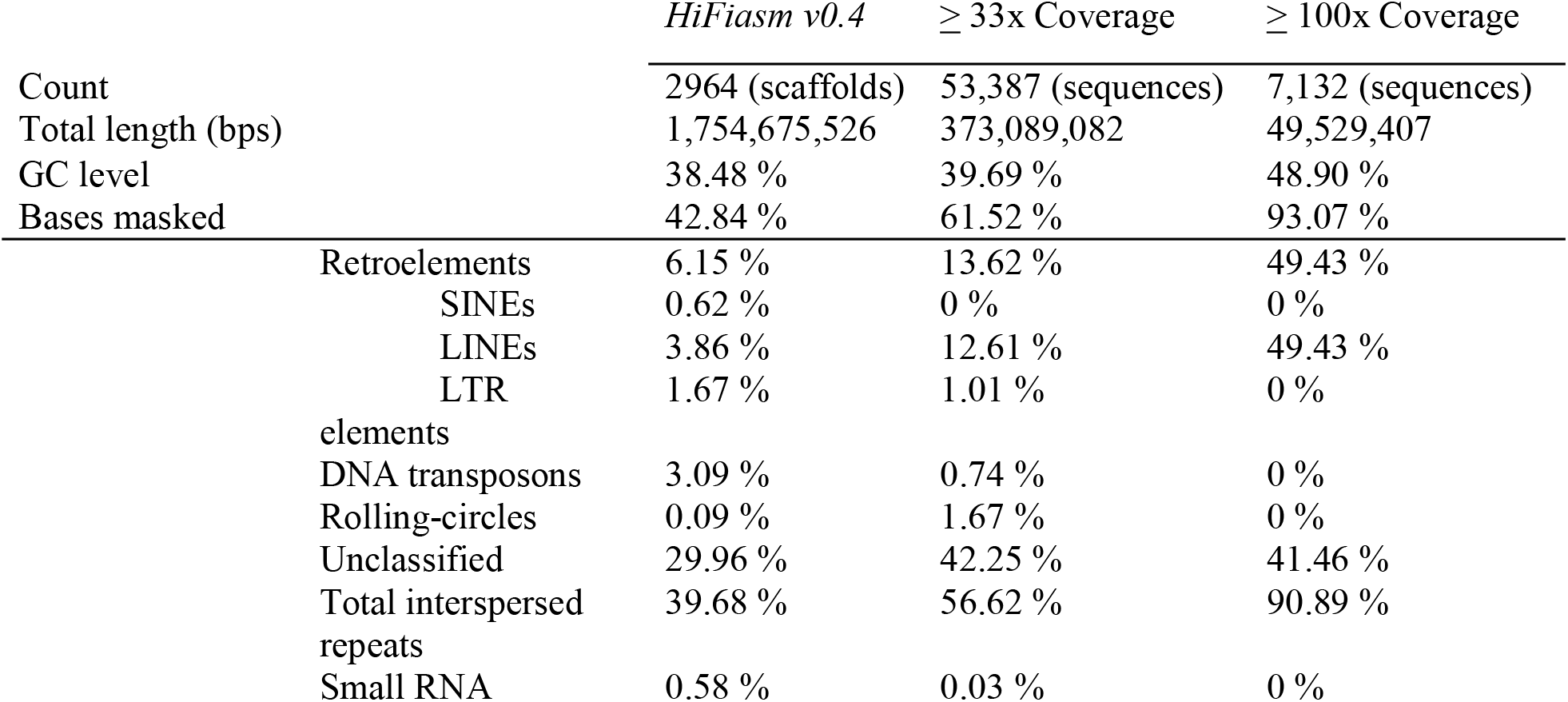
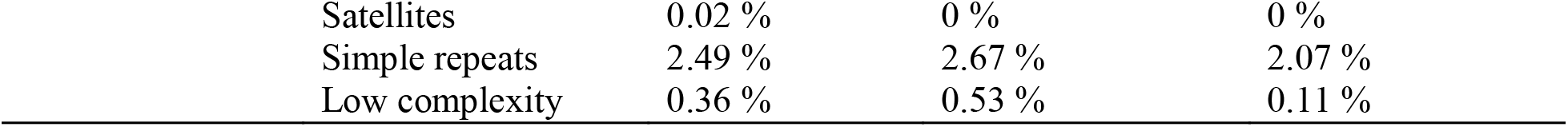
RepeatMasker results of the final assembly (*HiFiasm v0*.*4*), as well as additional read files that were extracted after raw reads were aligned back to the assembly, which represent reads with ≥ 33x coverage or ≥ 100x coverage of the genome. A custom *Diadema antillarum* repeat database was used for this analysis of repeats.

The Purge_Dups tool generated a ‘cleaned’ assembly file containing 1.15 Gb of data that removed the redundant contigs from the *HiFiasm v0*.*4* assembly, which was comprised of 1,235,546,339 bps (1.23 Gbps) and 1,024 contigs. This ‘cleaned’ assembly retained 70.41% of the bps in our *HiFiasm v0*.*4* assembly. The second output file contained 2,015 contigs across 519,138,994 bps (519 Mbps), with a file size of 495.2 Mb. This file contained the redundant contigs and had removed 29.59% of the bps from the *HiFiasm v0*.*4* assembly. The Purge_Dups function created additional contigs, which can happen when the algorithm splits a single contig into multiple contigs, extracts multiple regions of a single contig, or combines results from different alignment stages (Guan et al., 2020). Nonetheless, these results indicate that there was ∼2x as many contigs in the redundant file vs. the ‘cleaned’ assembly. In addition, the python output statistics reported that 99.36% of the redundant contigs had been removed from the ‘cleaned’ assembly. Lastly, the results of the BUSCO analysis on the ‘cleaned’ assembly (metazoan_odb10; n=954) indicated that it was 97.4% complete, containing 62.5% complete and single-copy BUSCOs, 34.9% complete and duplicated BUSCOs, 1.7% fragmented BUSCOs, and 0.9% missing BUSCOs.

The NCBI annotated *HiFiasm v0*.*4* assembly resulted in 33,123 protein-coding genes. In addition, there were 3,091 non-coding transcripts and 1,544 non-transcribed pseudogenes. The annotation (GCF_040938485.1-RS_2025_02) and features are available at https://www.ncbi.nlm.nih.gov/datasets/genome/GCF_040938485.1.

## Discussion

We present the first *de novo* genome assembly of *D. antillarum*. Our analysis pipeline utilized high-quality HiFi reads to generate a near complete assembly (98.4%), with key metrics supporting its completeness, contiguity, and overall robustness. We generated relatively large contigs, with a maximum length of 14.3 Mbp and an N50 of 1.56 Mbps, which suggests high contiguity. The per-base coverage plot shows that most of the genome is covered at a depth of 11x, with minimal missing regions, indicating the assembly accurately represents the genome. Additionally, the raw sequencing data used for assembly had an average quality score of Q34 or better, supporting the reliability of the genome assembly. The Blobtoolkit-generated snail plot provides further evidence of the high-quality assembly, with large contig sizes and high GC content (38.5%), which suggests a well-assembled and accurately represented genome.

Several features of the assembly quality metrics as well as the output statistics suggest the presence of duplications, repeat-rich regions, and heterozygosity in the genome. For instance, the BUSCO results suggest that most of the highly conserved housekeeping genes across metazoans have been duplicated (84.4%) in *D. antillarum*, and the Quast duplication ratio (2.52x) implies partial or whole genome duplication of the genome. Thus, we performed additional analysis to further characterize the extent of these features. We also addressed the presence of multiple haplotypes or otherwise pseudo-haplotypes in our assembly.

Firstly, we found no contamination of the haploid DNA sample used for the assembly either with DNA from another diploid species or prokaryote. In addition, if the snail plot and coverage plot had depicted two distinct peaks, then this would signify the presence of mixed haplotypes, pseudo-haplotypes, or rather a whole-genome duplication event (Challis et al., 2020; Ranallo-Benavidez et al., 2020; Vurture et al., 2017). While these plots lack the presence of two distinct peaks, there is a minor second bump present in the snail plot at ∼25x coverage. In addition, the coverage plot depicts a long tail that extends beyond the major peak at 11x coverage. These plot patterns suggest large-scale and/or numerous duplication events in the genome (Challis et al., 2020). This is further supported by the additional peaks in the Smudgeplots, which supports localized copy number variation (CNV) and recent segmental duplications. While the overall Smudgeplot pattern is diploid, certain regions of each plot show highly similar but heterozygous duplications in the genome (Ranallo-Benavidez et al., 2020).

There is large-scale duplication in the *D. antillarum* assembled genome based on the large structural variant analysis of reads with ≥ 1000 bps, but this includes only 0.31% of the genome, spanning 5.4 Mbps. A larger proportion of the repeat elements in the assembly, namely 30.53%, was extrapolated from targeting reads with ≥ 33x coverage of any bp length. This is nearly the same percentage of duplicated bps (29.59%) that were purged from the assembly using the purged_dups tool. In addition, ∼66% of the assembly contains redundant contigs (2,015 vs. 1,024). Lastly, the mapping of raw reads back to the assembly resulted in 0.38% of redundant mapping, which is also an indicator of genome duplication. While the coverage plot depicts that most of the genome is covered at 11x, there is extensive coverage beyond that (up to 530x), which is a sign of true duplicated reads as they are represented many times in the genome. Overall, this data suggests that approximately 30% of the genome has duplicated at some time in the past.

We reviewed the results of several analyses to address if the duplications in the assembly are real, or rather a product of collapsed reads, as shown by the small percentage of redundant read mapping (0.38%) of raw reads back to the assembly (Li, 2018; Vurture et al., 2017). Besides using long reads to assemble the genome, the coverage plot of the large structural duplications (≥ 1000 bps) mapped back to the assembly maintains consistency in coverage, suggesting that these regions accurately reflect the presence of multiple copies of sequences in the genome (see Supplementary Figure 3B). In addition, there is no evidence of collapsed reads in the Bandage plots (see Supplementary Figure 2), as these would be evident in contigs with higher coverage than 1x, which suggests that multiple copies had collapsed into a single node (Wick et al., 2015). Taken together, these results show that our assembly accurately reflects the duplication events that have occurred in the species. Some of the duplications in the genome may be composed of repeat elements, which comprise 42.84% of the *D. antillarum* genome. However, more research is needed to further classify the type of duplications in the genome, and in what regions of the genome these have occurred. In addition, there is a moderate level of heterozygosity (2.52%) in the genome. While the genome was assembled using a haploid sperm sample, the analysis of the unassembled HiFi reads suggests that heterozygosity is not so high as to cause ambiguity in the assembly process. The presence of duplicated regions and the higher coverage of some regions could be related to allelic variation, which is supported by this moderate level of heterozygosity in the genome.

Overall, the results of this study indicate that the *D. antillarum* genome harbors duplication, repeat elements, and heterozygosity. These features likely contribute to the complexity observed in the assembly, but they do not hinder the accuracy of the overall completeness or quality of the assembly. This assembly will serve as a valuable resource for future studies in comparative immunology, evolution and population genetics. In addition, it will provide a framework for the monitoring and restoration initiatives for this keystone marine invertebrate.

## Acknowledgments

Collection permits for animals include the DRNA permit number O-VS-PVS15-AG-00046-01082018 for An #1 and An #2, and the DRNA permit number 2021-IC-092 for An #3. The authors would like to thank Heidi D. Morales Díaz for assistance with the animal collections in Rincón. We also extend gratitude to the NCBI EGAP team for their expedited annotation of the GCA_040938485.1 assembly.

## Data Availability

Data generated for this study are available under NCBI BioProject PRJNA962153 and PRJNA839760. Raw ONT sequences for An #1 (NCBI BioSample SAMN46982352) generated 3.9 Gb of data containing 4.3 Gbps and An #2 (NCBI BioSample SAMN46998589) generated 2.2 Gb of data comprising 2.6 Gbps, which are deposited in the NCBI SRA under SRR32463885 and SRR32479314, respectively. Raw sequences for An #3 include PacBio HiFi reads (NCBI BioSample SAMN34386196) and Illumina MiSeq reads (NCBI BioSample SAMN34386197) that are deposited in the NCBI SRA database under SRR32365430 and SRR32478703, respectively. Additional NCBI GenBank accessions include the genome assembly (GCA_040938485.1), genome annotation (GCF_040938485.1-RS_2025_02), assembly reads (JBFRCO010000001 – JBFRCO010002964), and custom repeat database (SRR32898058).

## Author Contributions

A.J.M., J.M.W., T.K.O. and N.V.S. conceptualized the study. J.M.W., J.E.L., A.J.M. and T.K.O. acquired funds for the study. J.M.W., A.M.D.L. and A.J.M.C. conducted animal and sample collection. A.J.M.C., S.O.C.M., K.H., A.J.M. and J.M.W. extracted DNA. J.M.W. shipped samples for PacBio and Illumina sequencing. S.O.C.M., K.H., and A.J.M. generated sequencing libraries and conducted ONT sequencing. C.F.P. assembled the genome. A.J.M. and W.W. performed formal analysis on the assembled genome. A.J.M. drafted the manuscript. J.M.W., A.M.D.L., and N.V.S. edited the manuscript. All authors approved the final manuscript.

---

Supplementary Figure 1. Ploidy analysis of the adapter removed HiFi reads using Smudgeplot (v0.2.6). K-mer of size 21 (A) or size 31 (B) was used to infer ploidy in the *Diadema antillarum* genome. The plots display the distribution of normalized minor allele frequency (B / (A + B), x-axis) versus total k-mer pair coverage (A + B, y-axis), with color intensity representing the number of k-mer pairs. In (A), the observed genotype configurations, including AB, AAB, AAAB, and AABB have the respective frequencies (0.61, 0.23, 0.03, and 0.05). This indicates that there is an equal contribution of alleles for AB, while the proportion of AAB suggests segmental duplication or potentially a triploid like genome. The smaller ratios of AAAB and AABB suggest some presence of these additional alleles in the genome. In (B), the observed genotype configurations are AB and AAB, with frequencies of 0.73 and 0.18, respectively. The larger ratio of AB suggests a strong diploid signal, while the smaller proportion of AAB indicates some presence of additional alleles, or otherwise heterozygosity in the genome. The predominance of the AB genotype in both (A) and (B) plots indicate a diploid genome structure, although the presence of minor peaks corresponds to higher allele dosages, which suggests localized copy number variation or recent duplications in the genome (Ranallo-Benavidez et al., 2020).

Supplementary Figure 2. Quality assessment of the *HiFiasm v0*.*4* genome assembly. Bandage plots of the HiFi long-read genome assembly. Each plot shows the same 2,964 contigs. While plot (A) is unlabeled, plot (B) includes the coverage of each contig, which is 1x. Each contig is depicted as a curved line, with no overlap between structures. The longest contig in each plot is 14,326,224 bps long, and the shortest contig is 12,568 bps long. Base pair lengths have been omitted from these images to preserve clarity of the structures.

Supplementary Figure 3. Duplicate read characteristics in the *Diadema antillarum* genome. A. Histogram showing the distribution of read lengths for large duplicated structural variant (SV) reads (≥1,000 bp) that were mapped back to the genome assembly. The majority of these duplicated reads fall between 6,000 and 10,000 bp in length, suggesting long-read consistency within duplicated regions. The y-axis is plotted on a log scale to highlight both high- and low-frequency read lengths. B. Histogram of coverage depth across the genome for the same set of large duplicated SV reads. The height of each bar indicates the number of bases in the genome assembly that fall within a specific coverage range (e.g., 0–10x, 10–20x, etc.). Coverage depth extends up to 80x and displays a continuous distribution without sharp spikes, suggesting that the duplicated sequences are broadly and evenly distributed throughout the genome rather than clustered in specific hotspots. The y-axis is shown on a log scale to emphasize variation across several orders of magnitude.

