## Supplemental Table 1 for "A draft *de novo* assembly of *Diadema antillarum*, a keystone herbivore of the Caribbean reefs"

Supplementary Table 1. Assembly metrics compared for *D. antillarum* and *D. setosum*.

|  |  | *D. antillarum* | *D. setosum** |
| --- | --- | --- | --- |
| Sequencing platform |  | PacBio, ONT | PacBio, Omni-C |
|  | Genomescope^(Kmer 21)^ |  |  |
|  | Haploid genome length | 703 Mbps | 800 Mbps |
|  | Uniqueness (%) | 76.7 | 68 |
|  | Heterozygosity (%) | 2.52 | 2.11 |
| Assembler Type |  | hifiasm | hifiasm |
| BUSCO completeness (%) (metazoan_odb10) n=954 |  | 98.4 | 97.8 |
| No. assembled fragments (pseudochromosomes) |  | 2964 contigs | 101 (22) |
| Assembled genome size |  | 1.7 Gbps | 885.8 Mbps |
| GC (%) |  | 38.48 | 38.36 |
| Complete and single-copy BUSCOs (%) |  | 14.0 | 97.8 |
| Complete and duplicated BUSCOs (%) |  | 84.4 | 0.3 |
| No. protein coding genes |  | 33,123 | 23,030 |
| Characterized repeat elements (%) |  | 42.85 | 36.98 |
| Uncharacterized repeat elements (%) |  | 29.96 | 25.87 |

*Data obtained from Hong Kong Biodiversity Genomics Consortium et al. 2024.
