## Supplementary figures and images for "A draft *de novo* assembly of *Diadema antillarum*, a keystone herbivore of the Caribbean reefs"

### Supplemental Figure 1

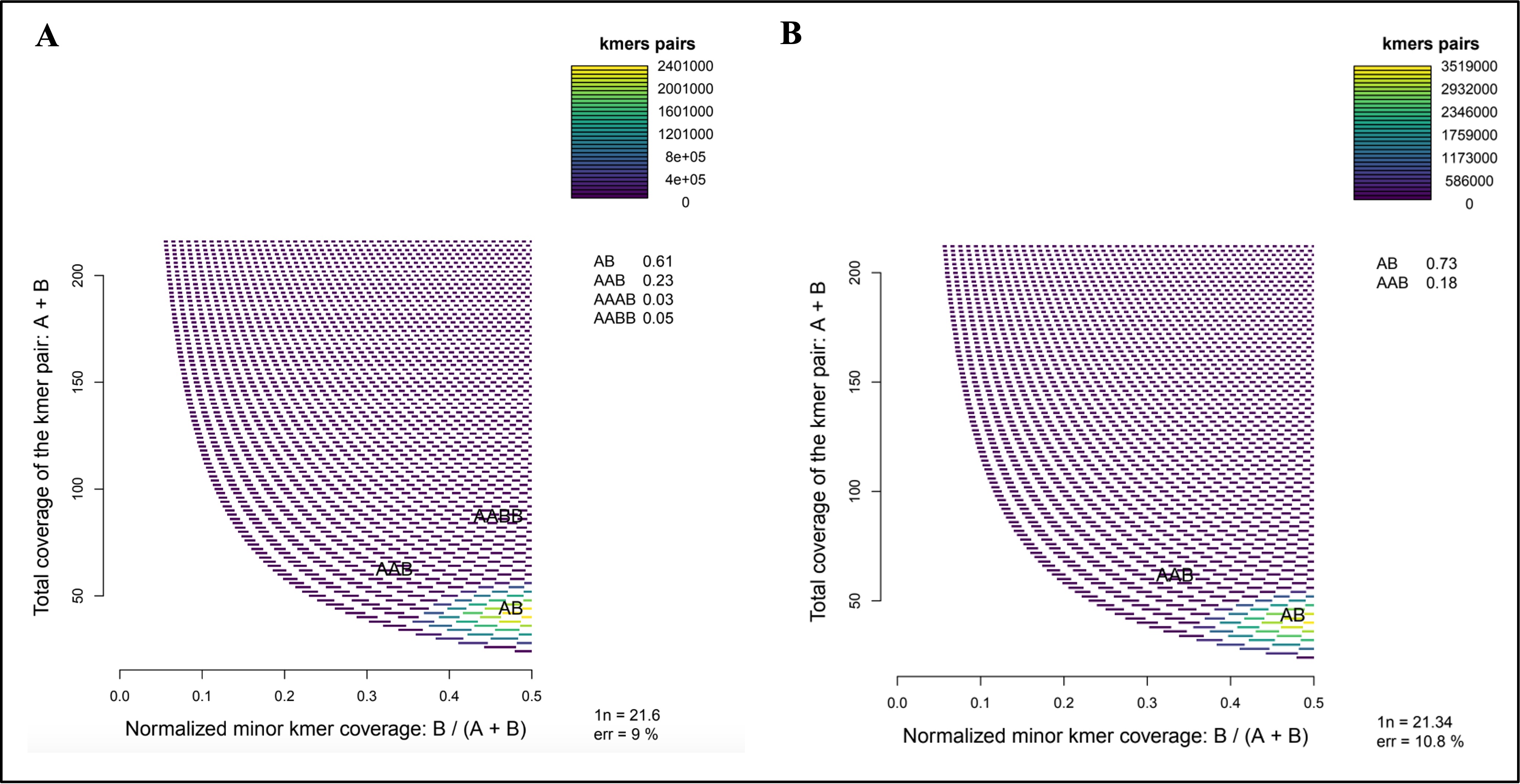

### Supplemental Figure 2

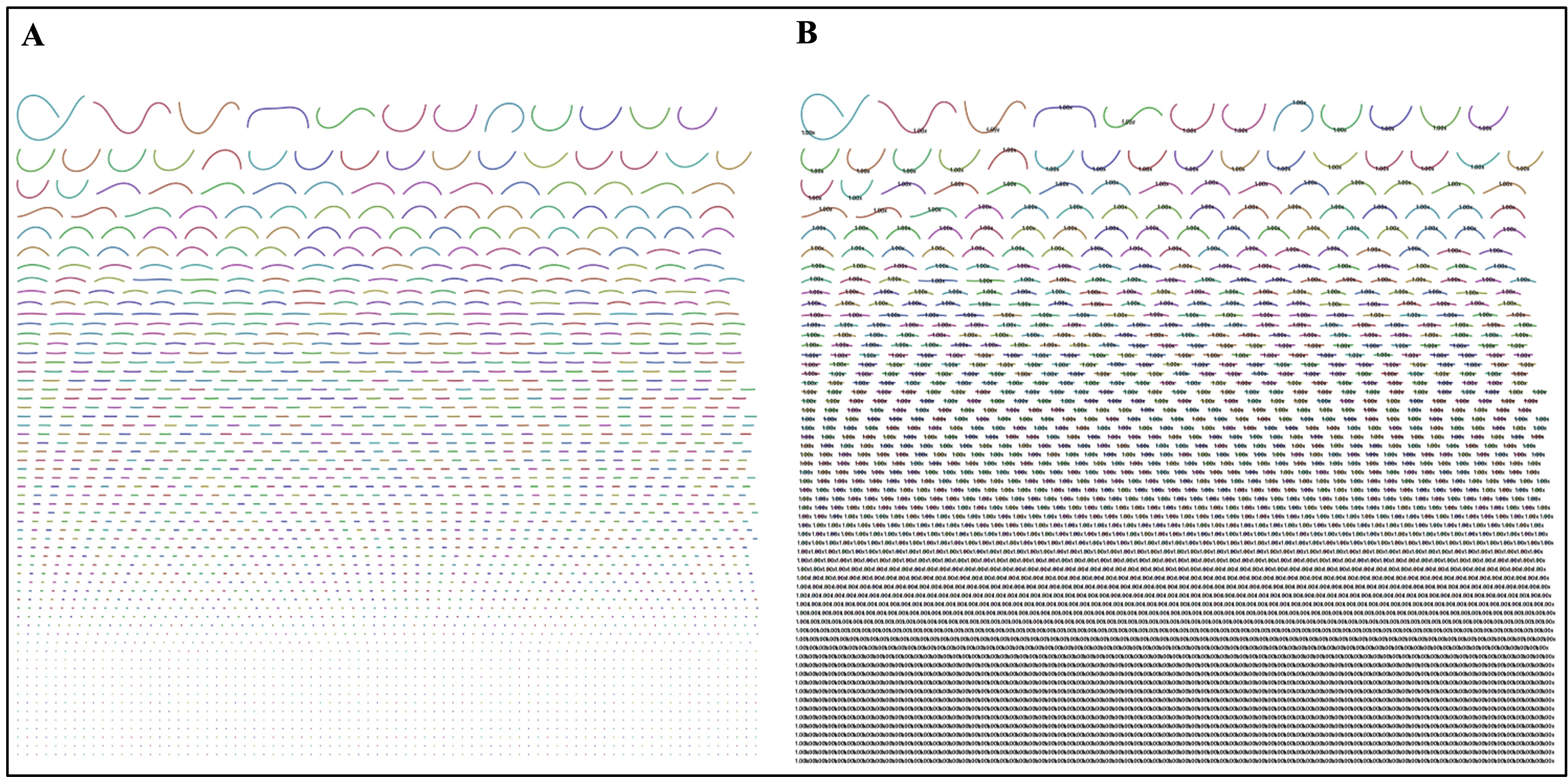

### Supplemental Figure 3

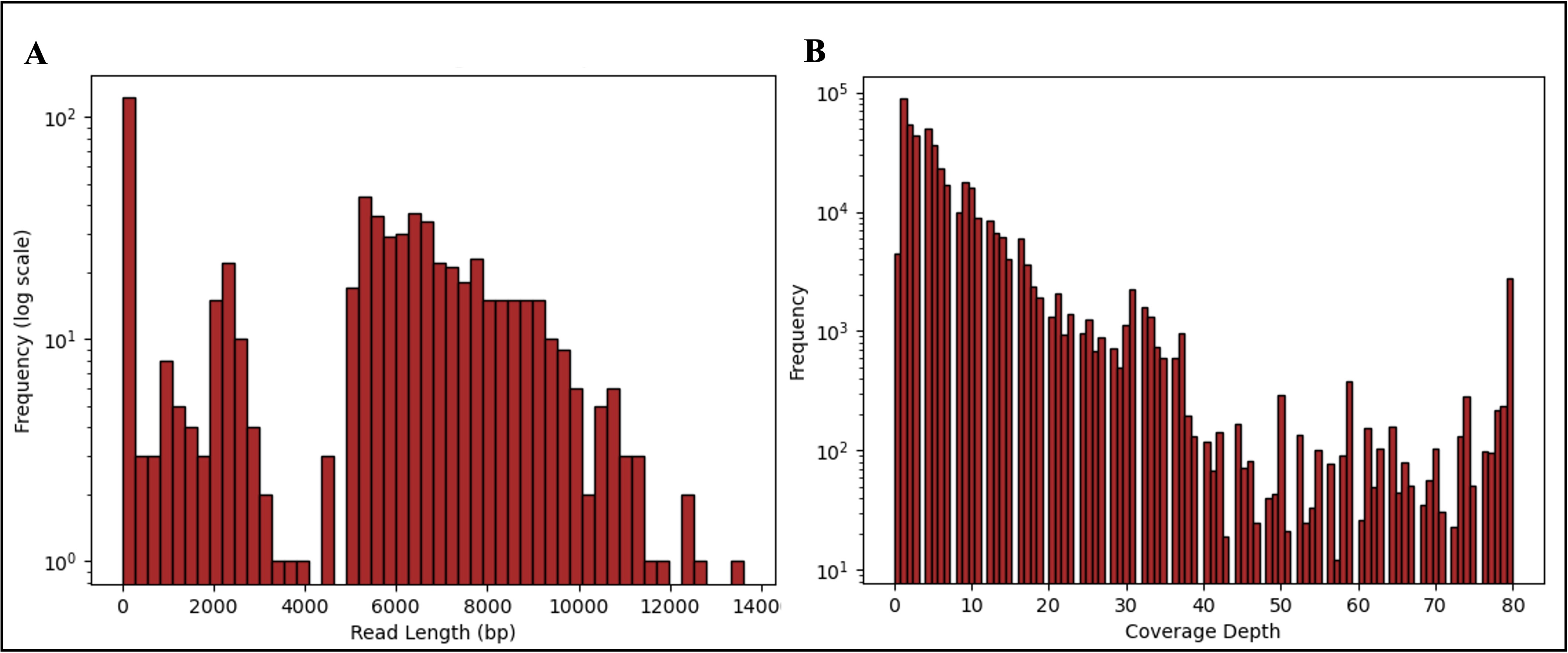
